# Ancient individuals from the North American Northwest Coast reveal 10,000 years of regional genetic continuity

**DOI:** 10.1101/093468

**Authors:** John Lindo, Alessandro Achilli, Ugo Perego, David Archer, Cristina Valdiosera, Barbara Petzelt, Joycelynn Mitchell, Rosita Worl, E. James Dixon, Terence E. Fifield, Morten Rasmussen, Eske Willerslev, Jerome S. Cybulski, Brian M. Kemp, Michael DeGiorgio, Ripan S. Malhi

## Abstract

Recent genome-wide studies of both ancient and modern indigenous people of the Americas have shed light on the demographic processes involved during the first peopling. The Pacific northwest coast proves an intriguing focus for these studies due to its association with coastal migration models and genetic ancestral patterns that are difficult to reconcile with modern DNA alone. Here we report the genome-wide sequence of an ancient individual known as *“Shuká <u>K</u>áa”* (“Man Ahead of Us”) recovered from the On Your Knees Cave (OYKC) in southeastern Alaska (archaeological site 49-PET-408). The human remains date to approximately 10,300 cal years before present (BP). We also analyze low coverage genomes of three more recent individuals from the nearby coast of British Columbia dating from approximately 6075 to 1750 cal years BP. From the resulting time series of genetic data, we show that the Pacific Northwest Coast exhibits genetic continuity for at least the past 10,300 cal BP. We also infer that population structure existed in the late Pleistocene of North America with *Shuká <u>K</u>áa* on a different ancestral line compared to other North American individuals (i.e., Anzick-1 and Kennewick Man) from the late Pleistocene or early Holocene. Despite regional shifts in mitochondrial DNA haplogroups we conclude from individuals sampled through time that people of the northern Northwest Coast belong to an early genetic lineage that may stem from a late Pleistocene coastal migration into the Americas.

**Significance Statement:** The peopling of the Americas has been examined on the continental level with the aid of single-nucleotide polymorphism arrays, next generation sequencing, and advancements in ancient DNA, all of which have helped elucidate major population movements. Regional paleogenomic studies, however, have received less attention and may reveal a more nuanced demographic history. Here we present genomewide sequences of individuals from the northern Northwest Coast covering a time span of ~10,000 years and show that continental patterns of demography do not necessarily apply on the regional level. In comparison with existing paleogenomic data, we demonstrate that geographically linked population samples from the Northwest Coast exhibit an early ancestral lineage and find that population structure existed among Native North American groups as early as the late Pleistocene.

## Introduction

The initial peopling of the Northwest Coast has received much attention due its proximity to Beringia and associated implications for an initial coastal migration into the Americas (1–3). Genetic clues for the peopling of the Northwest Coast, however, may be obscured by later demographic events in the region. Studies based on mitochondrial DNA (mtDNA) and Y-chromosomal markers suggest that populations in the region likely experienced admixture from other populations that entered the region after the initial peopling (4–6). Studies utilizing genome-wide data (7–9) inferred ancient gene flow into North America likely stemming from subsequent movements after the initial settlement. However, due to the limited genomic data from populations in this geographic region, those studies leave questions regarding the degree of temporal genetic continuity of Northwest Coast populations.

The distribution of the mtDNA haplogroup D4h3a in the Americas is also significant for the Northwest Coast region. Previous work on ancient individuals from North America revealed an interesting distribution of this haplogroup through space and time (Fig. 1). The oldest thus far sampled individual carrying this haplogroup is Anzick-1, dating back to ~12,600 cal BP and reportedly associated with Clovis technology (10, 11). Anzick-1 has proved to be surprising in a broader genetic sense, showing greater affinity with Central and South American groups than with Northern groups despite the burial being located in North America (but comparative indigenous populations from the United States are currently lacking). The same mtDNA haplogroup was originally identified on Prince of Wales Island, Alaska, in the individual named *Shuká <u>K</u>áa* (12). *Shuká <u>K</u>áa*, unearthed from On Your Knees Cave (OYKC), is not associated with Clovis culture but instead with a maritime tradition consistent with a coastal migration model and has been dated at ~10,300 cal BP (3). Approximately 300 km southeast of the OYKC archaeological site is Lucy Island, off the coast of British Columbia, Canada. This is the location of another individual bearing mtDNA haplogroup D4h3a, cataloged as 939, who died approximately 6,075 cal BP (13). Individual 939 displays genetic affinity to Pacific Northwest coast groups such as the Coast Tsimshian (henceforth "Tsimshian") who currently live in the same region but it is difficult to reject 939 as ancestral to both North and South American groups (9). This mtDNA haplogroup then nearly disappears from the area (or is found in extremely low frequency) in later ancient and living individuals (13–16). Less than 500 years after the time of 939, another previously published genetic sample (938 at ~5,675 cal BP), from an individual (13) found in the same Lucy Island archaeological site, exhibits the A2ag haplogroup. Notwithstanding the small number of samples analyzed from these time periods, the observed change in mitogenome haplogroups might be the result of gene flow into the region during the mid-Holocene. However, it is also possible that the mtDNA haplogroup A2 was already segregating in the population, and that the occurrence of the two mtDNA haplogroups at two different points in time may be explained by genetic drift or sampling.

**Figure 1.**
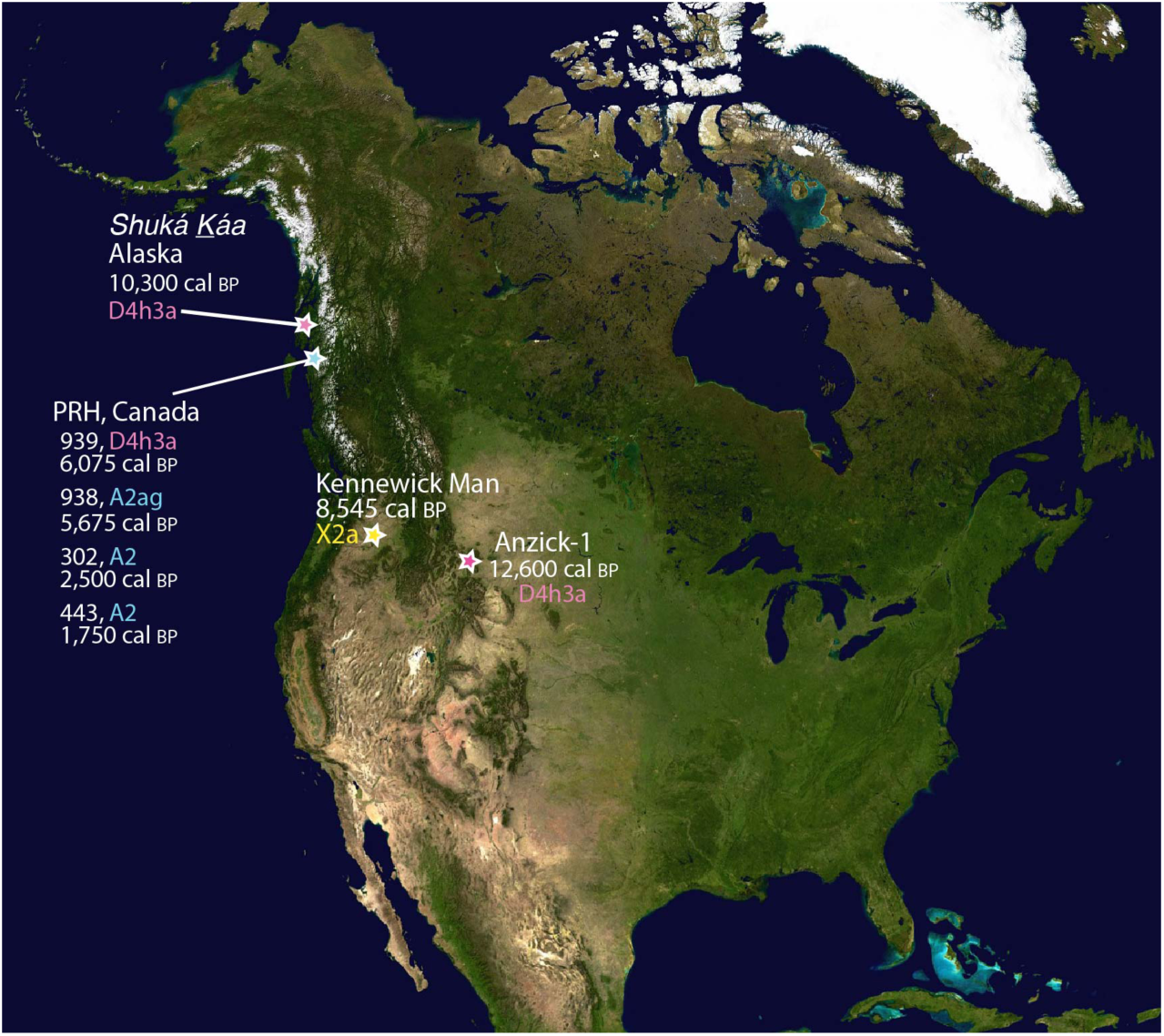
Sampling locations of ancient samples and their associated mtDNA haplogroups. The Anzick-1, *Shuká <u>K</u>áa*, and 939 samples all share the D4h3a mitochondrial haplogroup, whereas the 938, 302, and 443 samples from the Prince Rupert Harbour (PRH) region show a shift in mtDNA haplogroups to A2.

The only other ancient genome from North America is the Ancient One (also known as Kennewick Man), unearthed in the U.S. state of Washington and dating back to ~8,545 cal BP (17). Kennewick Man also displays surprising results as an early Holocene individual who resided in the Pacific Northwest. His mtDNA belongs to the northern North America limited haplogroup X2a, but his nuclear genome shows affinities with Central and South American populations, similar to patterns observed for Anzick-1, although a direct ancestry test demonstrates the greatest link to living individuals from the Confederated Tribes of the Colville Reservation, a Native population living in the same geographic region as Kennewick Man (17). On a broader scale, numerous areas of the Americas exhibit patterns consistent with genetic continuity of peoples in the same geographic region over time (9).

To test hypotheses related to different demographic scenarios for the peopling of the Northwest Coast, we generated genome-wide data (including the complete mitogenome) for *Shuká <u>K</u>áa* from Alaska (Table S1; Fig. 1). In addition, we generated two ancient, low coverage genomes, 302 and 443, from Prince Rupert Harbour (PRH), British Columbia (Table S1; Fig. 1), dating to 2,500 and 1,750 cal BP, respectively. Along with previously described genomes from the Americas, we test two hypotheses about the peopling of the Northwest Coast. We first test whether the people of this geographic region demonstrate temporal genetic continuity dating back to at least 10,300 cal BP. Second, we test whether the ancestors of the Northwest Coast experienced additional gene flow in the mid Holocene to further explore the previously observed shift in mtDNA haplogroups on the Northwest Coast.

### Community engagement

It is important to note that the interactions between scientists and indigenous community members associated with this study were and continue to be respectful. *Shuká <u>K</u>áa* is the indigenous name given to the ancient individual found in On Your Knees Cave on Prince of Wales Island in Southeast Alaska. His remains were identified in 1996, the same year in which the Ancient One was unearthed from the banks of the Columbia River near Kennewick, Washington. However, unlike the antagonistic relationships that were to develop over the handling of the Ancient One’s remains (18), co-author Terry Fifield, a U.S. Forest Service archaeologist, and other researchers engaged with Tlingit and Haida speaking communities in Alaska and developed strong working relationships with community members. The scientists and tribes created a partnership where community members were involved in nearly all decisions and aspects of the research project (19). During conversations that took place in 1996, 2004, and later, the partnering indigenous communities supported scientific studies, as they believed it would show concordance with oral histories (Klawock Cooperative Association and Craig Community Association 1996 and 2007 and Sealaska Heritage Institute 2004). In 2007, *Shuká <u>K</u>áa* was returned to the indigenous communities after initial research demonstrated that he lived around 10,300 years before present (BP) and exhibited mtDNA and Y-chromosome haplogroups that are only found among Native Americans (20). *Shuká <u>K</u>áa* was reburied on September 25, 2008. With appropriate engagement and discussions, our analyses were conducted on the last remaining tissue subsampled from *Shuká <u>K</u>áa*’s molars for DNA analysis prior to his repatriation to the Tlingit and reburial in his ancestral land.

Further south, RSM and JSC established a partnership with the Metlakatla and Lax Kw'alaams First Nations in 2007 to aid in the study of the population histories of those communities. They are located in the Prince Rupert Harbour (PRH) region of British Columbia. As part of the active partnership, RSM and JSC visit the communities on a regular basis to develop research studies, discuss interpretations of results as well as manuscripts written for peer-review publication. The First Nations agreed to allow destructive DNA analysis of ancestral individuals recovered from archaeological sites in the region and currently housed at the Canadian Museum of History. This includes the individuals labeled 939, 443, and 302 that are analyzed in this study.

## Results

### A mitochondrial reassessment

The entire D4h3a mtDNA haplogroup of the *Shuká <u>K</u>áa* individual was compared with 51 modern D4h3 mitogenomes (Datasheet S1) and with the other two published ancient mtDNAs (939 and Anzick-1). The resulting tree clearly shows that Anzick-1 is ancestral to the entire D4h3a clade, whereas the ancient Northwest Coast mitogenomes belong to two different sub-branches known as D4h3a9 and D4h3a12, with the latter here defined for the first time and encompassing a modern sample of an individual currently living in Bolivia (Fig. 2, Datasheet S1).

**Figure 2.**
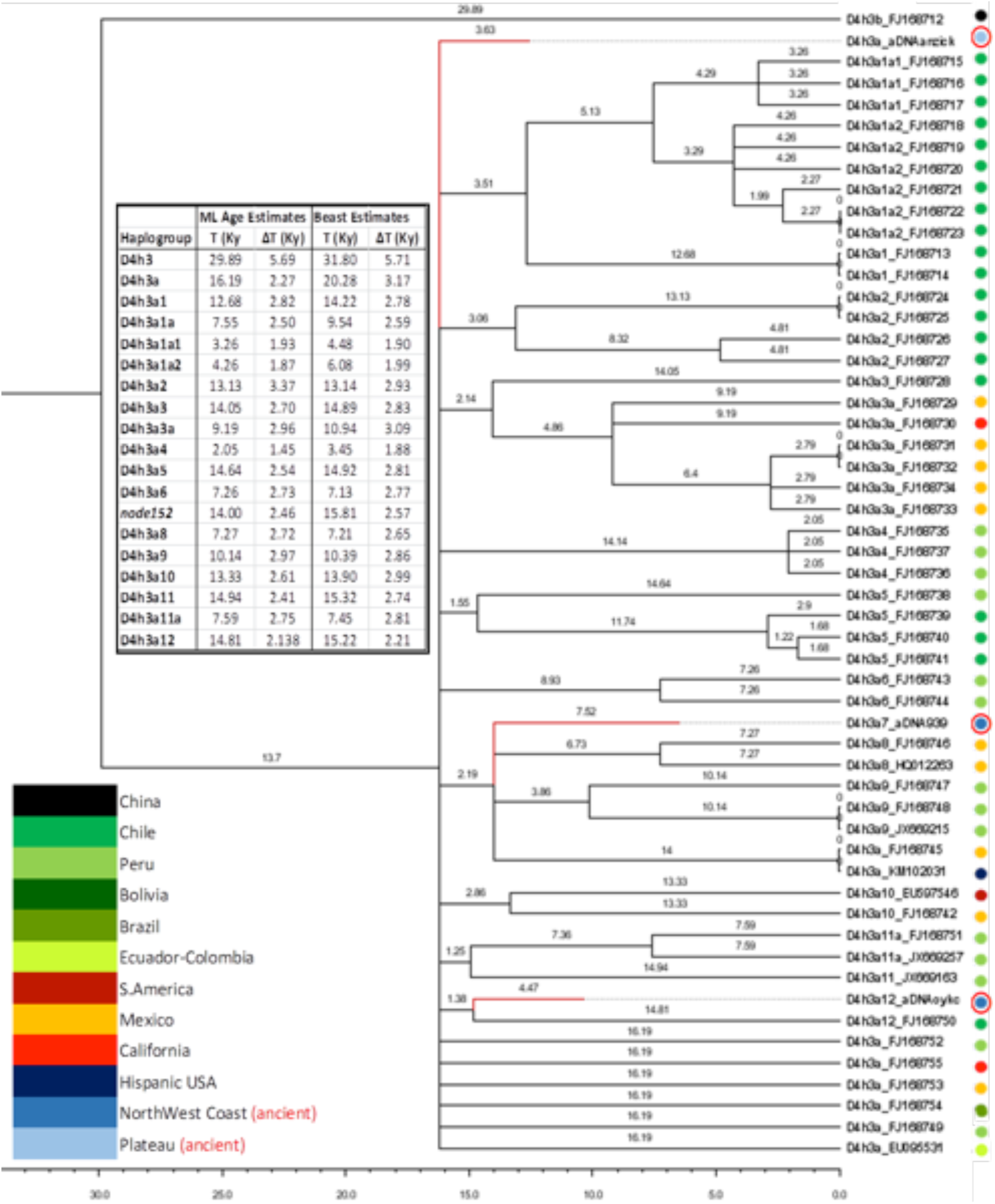
Phylogeny of mtDNA haplogroup D4h3. The topology was inferred by maximum parsimony. A maximum likelihood time scale is indicated below the tree. The insert shows the geographic distribution of major haplogroups based on these complete mtDNAs. Ancient mitogenomes are circled in red. Age estimates for the main nodes are in the inset.

It is worth noting that the 939 haplotype shares a mutation at nucleotide position 152 with two early South American branches D4h3a8 and D4h3a9. The radiocarbon dates of the three ancient mitogenomes were used as priors to date the most recent common ancestor between the Native American D4h3a and its sister Asian clade D4h3b at ~30–32 thousand years ago (kya). Soon after the Last Glacial Maximum the D4h3a branch differentiated (~16–20 kya) in northern North America possibly including the first settlers of the Northwest Coast. This ancestral group soon experienced a steep increase (more than 14x) of its effective population size while rapidly moving toward the Interior Plateau as attested by the Anzick-1 mitogenome, and southward, as testified by the ancient age of the D4h3a sub-clades currently found only in South America (Fig. 2, Datasheet S1).

Today, the haplogroup D4h3a is virtually absent in northern North America. To the contrary, the mitogenomes of the more recent ancients from the Northwest Coast (443 and 302) are classified as A2 (Table S1) the most commonly reported mitochondrial haplogroup of native North America. Thus, based on the mitogenome data alone it seems plausible that the native people of the northern Northwest Coast experienced a drastic change in their mtDNA gene pool in a rather short period of time possibly due to additional gene flow in the mid-Holocene (mitochondrial hypothesis). However, since mitochondrial DNA can describe only part of the ancestral genetic history of the Northwest we extended the analyses to the entire genome to test this hypothesis.

### Genome-wide assessment

We employed outgroup *ƒ*_3_ statistics to assess the shared ancestry between the ancient individuals and 169 worldwide populations (9). Outgroup *ƒ*_3_ statistics of a worldwide dataset demonstrate that all four ancient individuals (*Shuká <u>K</u>áa*, 939, 443, and 302) display greater affinity with Native American groups than with other worldwide populations (Figs. 3A, S2). Ranked outgroup *ƒ*_3_ statistics suggest that 939, 443, and 302 tend to share greatest affinity with Northwest Coast groups, whereas *Shuká <u>K</u>áa* ostensibly shows closer affinity to groups further south (Fig. S3). However, due to the low coverage of the *Shuká <u>K</u>áa* sample, the relationship is not statistically significant.

**Figure 3.**
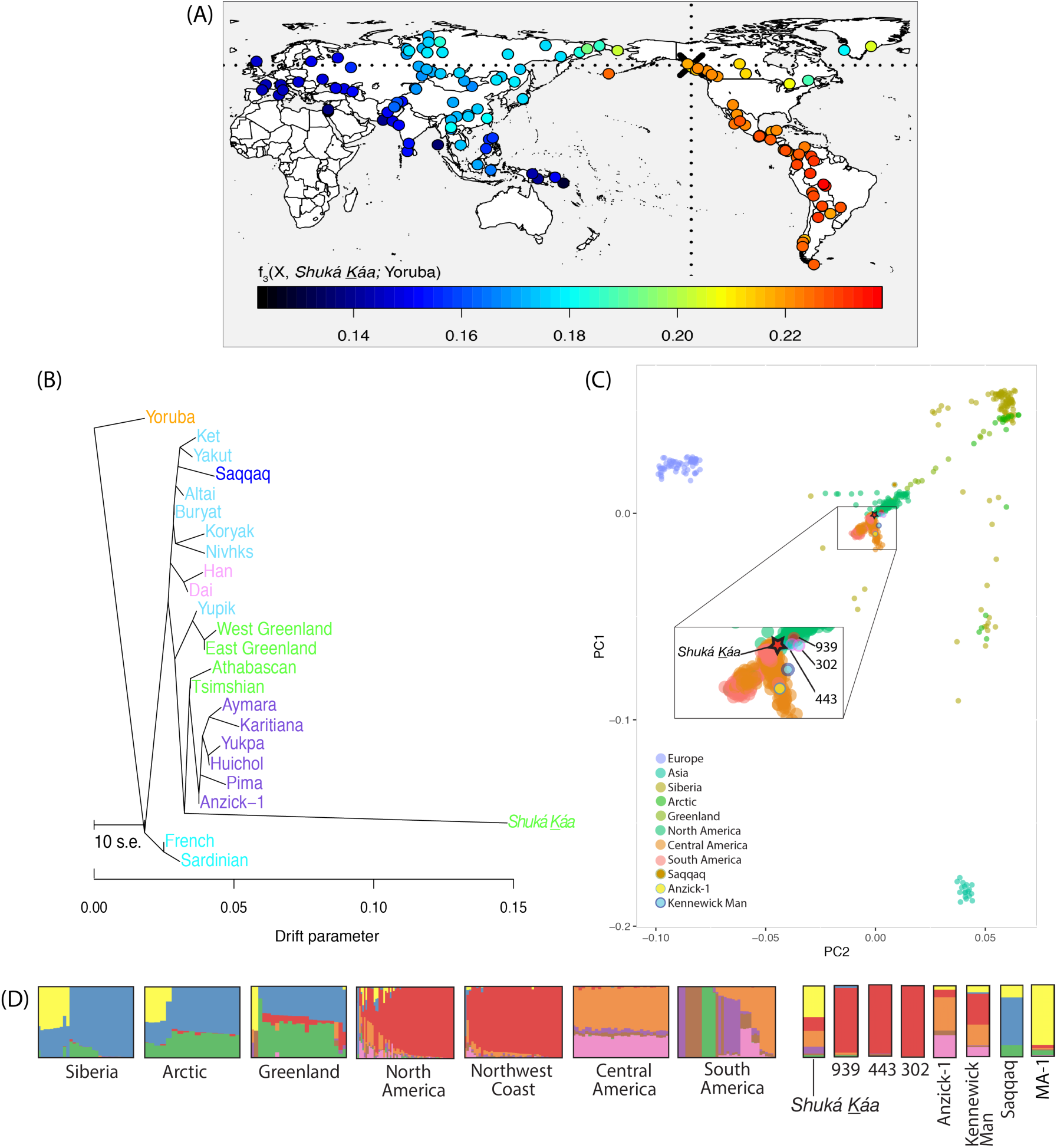
Genetic Affinity of *Shuká <u>K</u>áa* and the other Northwest Coast prehistoric humans to global and regional indigenous populations. (A) *Shuká <u>K</u>áa* shows greater genetic affinity with Native American groups than other global populations. Heat map represents the outgroup *ƒ*_3_ statistics estimating the amount of shared genetic drift between *Shuká <u>K</u>áa* and each of the 156 contemporary populations since their divergence with the African Yoruban population. (B) Maximum likelihood tree generated by *TreeMix* using whole-genome sequencing data from Raghavan et al. (9), and with the Tsimshian genome masked for European ancestry. (C) Principal components analysis projecting *Shuká <u>K</u>áa*, 939, 302, 443, Anzick-1(11), Saqqaq(21), and Kennewick Man (17) onto a set of non-African populations from Raghavan et al. (9), with Native American populations masked for non-native ancestry. (D) Cluster analysis generated by *ADMIXTURE* for set of indigenous populations from the Americas, Siberia, Artic, Greenland, and the Anzick-1, Kennewick, MA-1(22), Saqqaq, *Shuká <u>K</u>áa*, 939, 302, and 443 samples. The number of displayed clusters is *K*=8, which was found to have the best predictive accuracy given the lowest cross validation index value.

To further elucidate the relationship among the ancient individuals of the Northwest Coast and their relationship to modern populations, we examined maximum likelihood trees created with *TreeMix* (23). C/T and G/A polymorphic sites were removed from the data set to guard against the most common forms of post-mortem DNA damage (24). We observe that 302 and 443 form a sister clade to the modern Tsimshian (masked for European ancestry) (Figs. S4A and B, respectively). Individual 939 is an outgroup to both North and South Americans (Fig. S4C), as is *Shuká <u>K</u>áa* (Fig. 3B). However, adding a migration event introduces an edge connecting Europeans and *Shuká <u>K</u>áa* which leads to *Shuká <u>K</u>áa* forming a clade with the Tsimshian and Athabascan (Fig. S5). The signal may represent Native American dual ancestry (22) or be a result of possible contamination (Table S2).

Principal components analysis reveals a tight clustering of 939, 443, and 302 which also overlaps with modern North American indigenous populations (Figs. 3C, S6). *Shuká <u>K</u>áa* falls in close proximity but overlaps with both North and South American groups. The admixture clustering analysis shows a more complicated pattern where *Shuká <u>K</u>áa*, exhibits mainly components identified in North American and Siberian/Arctic individuals, as well as smaller fractions found in South American populations (Fig. 3D). However, individuals 939, 302, and 443 all exhibit a major component found in North American populations.

To further test the hypothesis that people of the Northwest Coast exhibit a close genetic relationship with ancient individuals from the same region, we employed the *D* statistic (25). Given the *TreeMix* admixture results between *Shuká <u>K</u>áa* and European populations we performed a contamination correction to the *D* statistic as described in Raghavan et al. (22) utilizing observed *D* statistics with European populations (*SI Appendix*). Hypothetical scenarios based on the *D* statistics are depicted in Figure 4. The *D* statistic does not support a scenario of genetic continuity between Anzick-1 and *Shuká <u>K</u>áa* with respect to South Americans (Fig. 4A, Table S3). The relationship of *Shuká <u>K</u>áa*, however, is more complex when examined with specific North American ancient and modern groups. Comparing *Shuká <u>K</u>áa* with 939 and the more recent ancient individuals from the Northwest Coast (443 and 302), we cannot reject an equally diverged relationship with respect to *Shuká <u>K</u>áa* (Fig. 4B, Table S3). However, we see the same relationship when the comparison is performed between both ancient or modern Northwest Coast individuals and individuals from South America where *Shuká <u>K</u>áa* is basal to both groups (Fig. 4C, Table S3). Individual 939 displays a similar pattern with *TreeMix* and *D* statistics (Fig. S4C, Table S3) where the individual appears basal to both the Northwest Coast and South America. However, the admixture results show a predominately “North American” component and contamination-corrected *D* statistics for 939 indicate a significant relationship toward the Northwest Coast (Table S3; Tests 17, 18).

**Figure 4.**
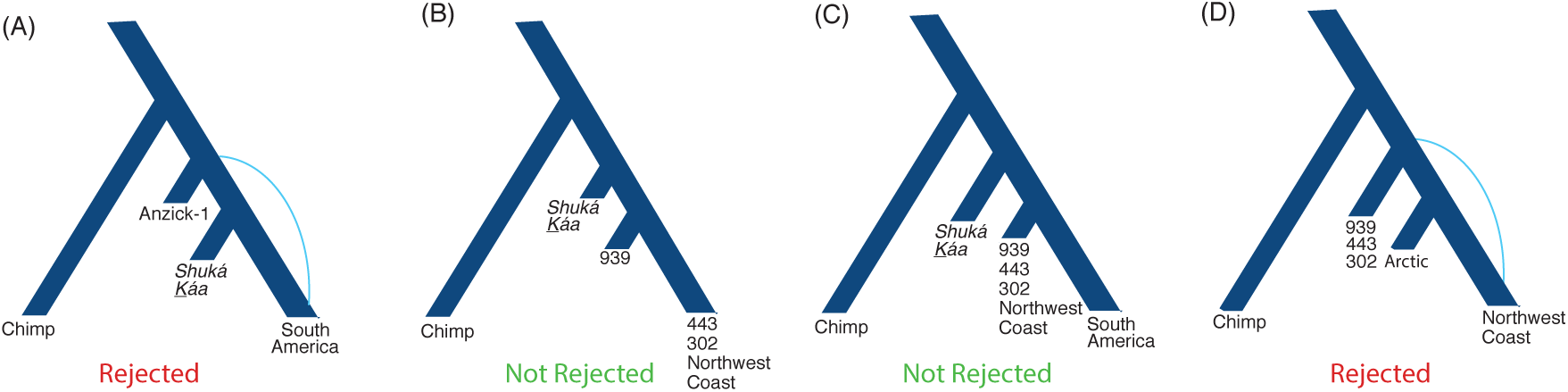
Hypothetical scenarios for the regional peopling of the Northwest Coast. We tested alternative models of shared ancestry of the ancient Northwest Coast individuals with contemporary and ancient populations of the Americas. Supporting *D* statistics are listed in Table S3. Additional sequenced samples were derived from Raghavan et al. (9). **(A)** Scenario tested by the *D* statistic where Anzick-1 is basal to both *Shuká <u>K</u>áa* and South America, which is rejected indicating a closer affinity to South America. **(B)** Scenario tested by the *D* statistic where *Shuká Káa* is basal to 939 and both contemporary and ancient Northwest Coast (NWC) individuals, which is not rejected. **(C)** Scenario tested by the *D* statistic where *Shuká <u>K</u>áa* is basal to ancient and modern NWC and South America, which is not rejected. **(D)** Scenario tested by the *D* statistic where 939, 443, and 302 are basal to arctic (Yup’ik and Inuit) and contemporary NWC populations, which is rejected indicating a closer affinity to contemporary NWC populations.

The *D* statistic did not reveal a signal of gene flow between Arctic populations (Inuit and Yup’ik) and either the modern or ancient Northwest Coast populations when compared to *Shuká <u>K</u>áa* (Table S3; Tests 19–28). However, when comparing the more recent ancient individuals, the tree was rejected with 939, 302, and 443 indicating greater affinity toward the Northwest Coast populations than to the Arctic (Fig. 4D, Table S3).

Because certain *D* (Table S3, Tests 8, 9) and *ƒ*_3_ (Fig. S3D) statistics yielded nonsignificant results, we next wanted to examine whether the basal relationship that *Shuká <u>K</u>áa* exhibited to Northwest Coast and South American populations could be due to its age relative to the split time of those groups. To address this hypothesis, we simulated genetic data with FastSimCoal2 (26), which allowed us to sample *Shuká <u>K</u>áa* 10,300 years in the past (*SI Appendix*). We considered one scenario (Scenario 1) in which *Shuká <u>K</u>áa* is on the branch leading to the Northwest Coast, and another scenario (Scenario 2) in which the sample was on a branch that diverged earlier than the split of the Northwest Coast and South American populations (Fig. S7A). Results for the 1000 simulated replicates under each scenario are plotted in Figure S7B indicating that only a small fraction of simulated replicates from Scenario 1 could reject the null hypothesis that *Shuká <u>K</u>áa* is equally diverged to the Northwest Coast and South American lineages even though the simulations placed *Shuká <u>K</u>áa* on the Northwest Coast branch. However, the reason for this lack of power may be due to the amount of data (which we controlled to yield a similar number of *D*-statistic informative sites as the empirical data). We therefore also considered a set of simulations where we increased the expected number of *D*statistic informative sites by an order of magnitude. Results from these simulations (Fig. S7C) show that the clear majority of simulated replicates from Scenario 1 could reject the null hypothesis that *Shuká <u>K</u>áa* is equally diverged to the Northwest Coast and South American lineages with *Shuká <u>K</u>áa* having higher affinity to the Northwest Coast. Further, results for Scenario 2 indicate that the null hypothesis is generally not rejected as expected from the simulated scenario (Figs. S7B, C).

We next considered models that examine a less direct relationship between *Shuká <u>K</u>áa* and the more recent ancient individuals in addition to the spread of the D4h3a mtDNA haplogroup across the Americas. Because Anzick-1 from Montana predates *Shuká <u>K</u>áa* (~12,600 vs. ~10,300 years BP) and shares the same mtDNA haplogroup we performed *D* statistics to assess their relationship. The tree with *Shuká <u>K</u>áa* and 939 as sisters relative to Anzick-1 is not rejected (Fig. S8A). The trees with modern Northwest Coast (mtDNA haplogroup A2) as sister to either 443 (mtDNA haplogroup A2) or 302 (mtDNA haplogroup A2) relative to 939 (mtDNA haplogroup D4h3a) are also not rejected (Fig. S8B). These results are inconsistent with a change in the overall gene pool of the Northwest Coast after the early Holocene even though it led to the observation of very different mitochondrial haplogroups detected over time in the region.

We also explored the relationship of *Shuká <u>K</u>áa* to Kennewick Man (17) who was unearthed in Washington and dates to ~8,545 cal years BP. Although Kennewick Man does not share the same mtDNA haplogroup with *Shuká <u>K</u>áa*, they co-existed within 1,700 years of each other (about 68 generations). In all tests with the *D* statistic, *Shuká <u>K</u>áa* displayed a basal relationship to both ancient and modern Northwest Coast populations with respect to Kennewick Man (Fig. S8C, Table S3).

## Discussion

These data support a shared ancestry for the indigenous peoples of the Northwest Coast, dating back to at least ~10,300 calendar years BP. The individual supporting this scenario is *Shuká <u>K</u>áa* who belonged to mtDNA haplogroup D4h3a. While both Anzick-1 and 939 also belonged to this mtDNA haplogroup later ancient and modern individuals of the Northwest Coast do not (13, 14). Despite belonging to different mtDNA haplogroups, *Shuká <u>K</u>áa* exhibits a close nuclear DNA relationship with 302 (~2,500 cal years BP) and 443 (~1750 cal years BP), both of whom belong to mtDNA haplogroup A2 which is observed today at high frequencies among modern Northwest Coast populations (14). Thus, based on mtDNA data alone, the hypothesis of a change in the genetic composition of the Northwest Coast appearing by the time of 302 and 443 seems plausible. However, our more extended genome-wide analysis does not support a change in the genetic composition of the Northwest Coast appearing by the time of 302 and 443, both of whom form a sister clade with the Tsimshian (Fig. S4). Instead, we observe a trend of genetic continuity through time, which is exemplified by individual 939 who displays affinities with both the more recent Northwest Coast ancient individuals and *Shuká <u>K</u>áa*. Individual 939 shares the same mtDNA haplogroup D4h3a as *Shuká <u>K</u>áa* while belonging to the predominant North American ancestry component observed in 443 and 302 (Fig. 3D). Furthermore, previously published data from individual 938 (13), who is similar in age and found on the same island site as 939, exhibits an A2 mtDNA haplogroup. These results indicate that the two haplogroups were already present in the ancestral population and are not the result of later gene flow into the area.

Also of interest is the evidence for the placement of *Shuká <u>K</u>áa* on different lineages than Anzick-1 and Kennewick Man (Fig. S8D). Despite their shared North American geographies and time periods as well as evidence that a single peopling event occurred in the late Pleistocene (27), our analyses demonstrate that population structure existed in the late Pleistocene of North America. These data are concordant with the late Pleistocene archaeological record documenting northward movement of the North American Paleo-Indian archaeological tradition following the establishment of a biotically viable deglaciation corridor between eastern Beringia and areas south of the merged Cordilleran and Laurentide ice sheets after 13,000-12,500 cal BP (28). These analyses further support archaeological data demonstrating that the contemporaneous population and archaeological structure is reflected in the late Pleistocene archaeological record.

We find that the placement of *Shuká <u>K</u>áa* (based on *D* statistic simulations) is consistent with residing on a branch with modern-day indigenous people from the Northwest Coast, whereas Anzick-1 fits separately on a branch that leads to the southern lineage, including populations from Central and South America (Fig. 5). Our result suggests that *Shuká <u>K</u>áa* was part of a population closely related to the ancestors that gave rise to the current populations of the northern Northwest Coast.

**Figure 5.**
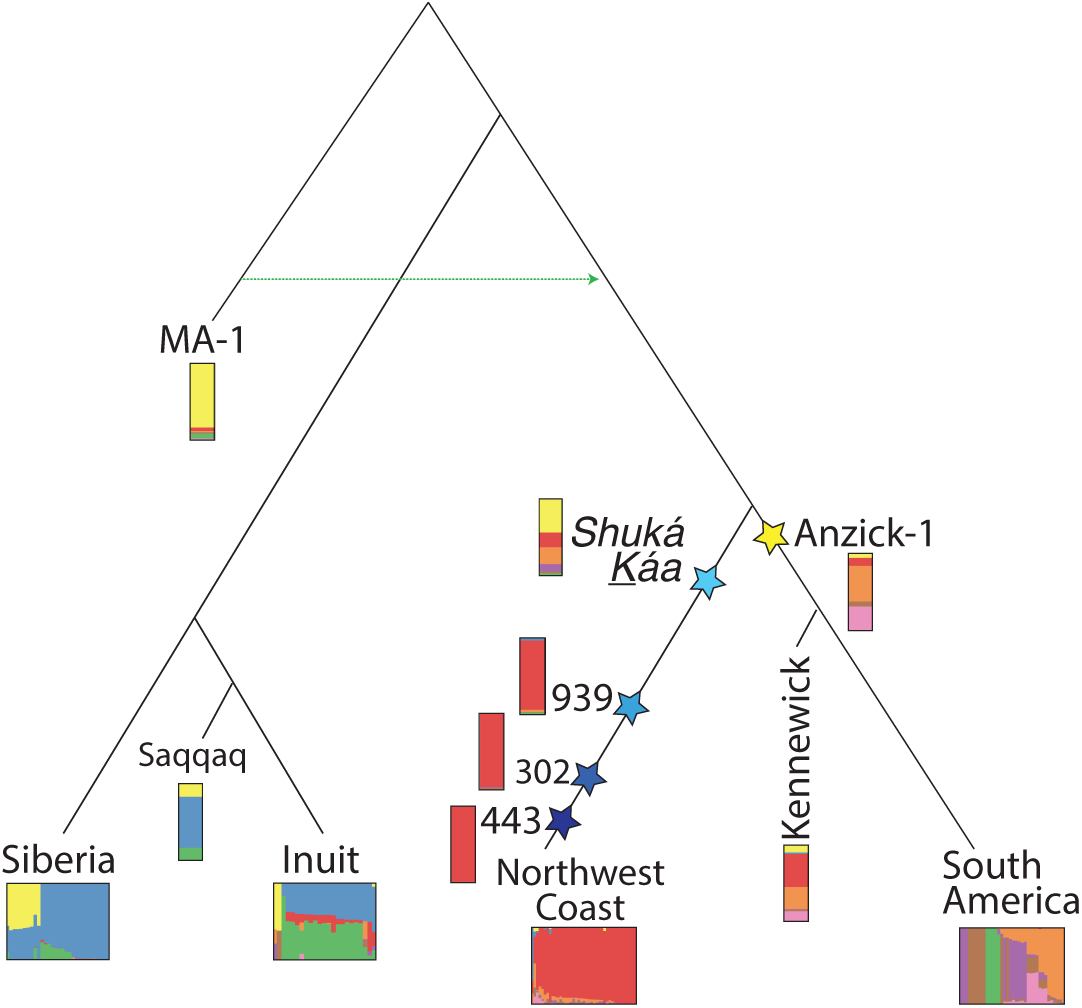
*Shuká <u>K</u>áa* in relation to other Native American groups. Schematic showing *Shuká <u>K</u>áa* placed on the branch leading to North Americans, which is supported by simulation-based *D* statistics.

On a broader scope, *ADMIXTURE* analysis revealed a component that dominates contemporary North American populations which increases over time regarding the Northwest Coast (Fig. 3D). We also observe a small fraction of this component in the 24,000 year-old MA-1 individual from Siberia (22) (Fig. 3D). This result suggests that the North American component reflects an early ancestral lineage that has drifted through time to high frequency on the Northwest Coast.

We conclude that the Northwest Coast exhibits an ancestral lineage that stems from the initial peopling of the region. The observed temporal change of mtDNA haplogroups in the area was probably due to sampling, and no clear signs of gene flow into the area after the first settlement have been identified. *Shuká <u>K</u>áa*, who lived some 10,300 years ago, was part of an ancestral population that may have first populated the region but was distinct from ancestral populations related to Anzick-1. Although we cannot use our data to identify the specific ancestral location of this coastal lineage, considering that *Shuká <u>K</u>áa* lived over 10,000 years ago and the interior corridor was not viable for human migration until after 12,600 years before present (29), it is unlikely that the lineage existed south of the North American ice sheets. The results presented here reveal the power of regional studies to elucidate demographic complexities, shedding light on the peopling of the Northwest Coast and the early ancestral lineages of North America.

## Methods

### DNA Extraction and Library Preparation

We used standard ancient DNA extraction methods following stringent guidelines to work with ancient human remains and conducted these in dedicated ancient DNA laboratories. DNA was extracted from 3 teeth belonging to individuals *Shuká <u>K</u>áa*, 302, and 443. Further, each DNA extract was converted into Illumina libraries (*SI Appendix*).

### Genome Enrichment

We captured from eight libraries of *Shuká <u>K</u>áa* DNA with the MyBaits whole genome enrichment kit, enhanced with protocol modifications recommended for the study of ancient DNA (*SI Appendix*). The eight captured libraries from each individual were pooled and sequenced (single-end) on four lanes of an Illumina HiSeq 2000 run. Two captured libraries from 302 and 443 were pooled and run on two additional lanes.

### Contamination Estimates

Contamination estimates utilizing the mitochondrial genome were run on all three samples, using the Scmutzi program described in Renaud et. al. (30). The method jointly estimates present-day human contamination and reconstructs the endogenous mitochondrial genome by considering both deamination patterns and fragment length distributions. Since *Shuká <u>K</u>áa* and 443 were typed as male, using the method described in Skoglund *et al.* (31), contamination based on the X chromosomes was also performed for these samples using the method described in Korneliussen *et al.* (32) and applied through the ANGSD software suite (http://popgen.dk/angsd). The *Shuká <u>K</u>áa* sample did not have sufficient coverage along the X chromosome to perform the estimate.

### *f*_3_ and *D* Statistics

To test the genetic affinity of the ancient individuals with global populations, we performed *f*_3_ outgroup statistics, using the method outlined by Patterson et al. (2012). We also examined the genetic affinities of each individual and various populations using ranked *f*_3_ statistics, shown in Figure S3. To examine the relationship between the ancient individuals (*Shuká <u>K</u>áa*, 939, 443, 302, and Anzick-1), we performed an ABBA-BABA test or *D* statistic (33) using the definition employed by ANGSD (32). The chimpanzee genome was used as an outgroup sequence. The tests also included the whole genome of a contemporary Tsimshian (9), which was masked for European ancestry, and all comparisons with *Shuká <u>K</u>áa* employed a correction for European contamination, as did several comparisons with 939 (*SI Appendix*). To guard against potential bias from DNA damage in the ancient individuals, transitions were not considered during the tests.

## Acknowledgments

This project was made possible through the active partnerships of the Lax Kw’alaams and Metlakatla First Nations and the Sealaska Heritage Institute. The field research was conducted in partnership with the Klawock Cooperative Association (Tribe) and the Craig Community Association (Tribe). The field work and preliminary analyses were supported by the National Science Foundation (OPP-99-04258, 97-22858, BCS-1413551 & BCS-1518026), the USFS Tongass National Forest, and the National Geographic Society. Research for this analysis was funded by the Office of the Vice Chancellor of Research, University of Illinois at Urbana-Champaign, by the Canadian Museum of History, Gatineau, Quebec, Canada, and by Pennsylvania State University startup funds. The ancient data have NCBI Short Read Archive accession no. _. Portions of this research were conducted with the Advanced CyberInfrastructure computational resources provided by The Institute for CyberScience at Pennsylvania State University. We thank Cara Monroe for assistance in the laboratory at Washington State University.

